# Super-Resolution Electrochemical Impedance Imaging with a 100 × 100 CMOS Sensor Array

**DOI:** 10.1101/2021.08.19.456729

**Authors:** Kangping Hu, Christopher E. Arcadia, Jacob K. Rosenstein

## Abstract

This paper presents a 100 × 100 super-resolution integrated sensor array for microscale electrochemical impedance spectroscopy (EIS) imaging. The system is implemented in 180 nm CMOS with 10 *μ*m × 10 *μ*m pixels. Rather than treating each electrode independently, the sensor is designed to measure the mutual capacitance between programmable sets of pixels. Multiple spatially-resolved measurements can then be computationally combined to produce super-resolution impedance images. Experimental measurements of sub-cellular permittivity distributions within single algae cells demonstrate the potential of this new approach.

## I. Introduction

Electrochemical impedance spectroscopy (EIS) is a powerful tool for chemical and biological sensing, with numerous applications in cell culture monitoring and biomolecular diagnostics [1], [2]. However, while biological samples have important spatial variation in their conductivity and dielectric properties, impedance is often recorded at only a single point in space. Some approaches add spatial dimensions using scanning probes [3] or small arrays of macroscale electrodes [4], but many opportunities remain to take advantage of the density and scale of CMOS integrated electrode arrays [5]–[7]. Integrated impedance imaging arrays can offer greater throughput than discrete electronics, faster acquisition than scanning probes, and finer spatial resolution than existing impedance tomography systems.

To address these challenges, we introduce a 100 × 100 CMOS EIS sensor array that uses an area-efficient two-phase switching scheme to measure the mutual capacitance between pairs of nearby pixels. The pixel grid pitch is 10 microns, and a measurement from the array can be considered as a radio-frequency impedance image which is a function of both the sensor parameters and the spatial distribution of dielectric properties within the sample above the sensor. The pixels are addressable in a pairwise manner. For example, we can record an image that represents the impedance between each electrode and the pixel one position to its left; we can then acquire a second image that describes the mutual capacitance between each pixel and the electrode two positions to its left.

By acquiring impedance images with different pairwise pixel offsets, we can assemble a high-resolution composite image from multiple frames of the same scene, using oversampling principles similar to those used for superresolution optical image reconstruction [8]. Compared to previous CMOS capacitance imaging arrays [5], [6], this new scheme only requires one extra pair of switches per pixel.

This paper is organized as follows. Section II explains the two-phase scheme and its measurement principle. In Section III we detail the system architecture and circuit design of a prototype EIS array implementation. Section IV presents experimental results in which we resolve sub-cellular features in green microalgae, and Section V concludes the paper.

## II. Two-Phase Mutual Capacitance Sensing

To illustrate the two-phase sensing scheme, Fig. 1(a) presents a model of two electrodes that are capacitively coupled to a buffer solution. *C*_1_ describes the capacitance that is only seen by electrode #1, and *C*_2_ is the capacitance that is only seen by electrode #2. *C_M_* is the mutual capacitance between these two electrodes, which may include distributed electric fields extending into the sample as well as parasitic capacitance within the sensor chip. We neglect the effects of Debye shielding because the circuit is operating at radio frequencies [5], [6], [9]. We also assume that the capacitors charge faster than the switching cycle so that we can neglect any distributed resistance.

**Fig. 1:**
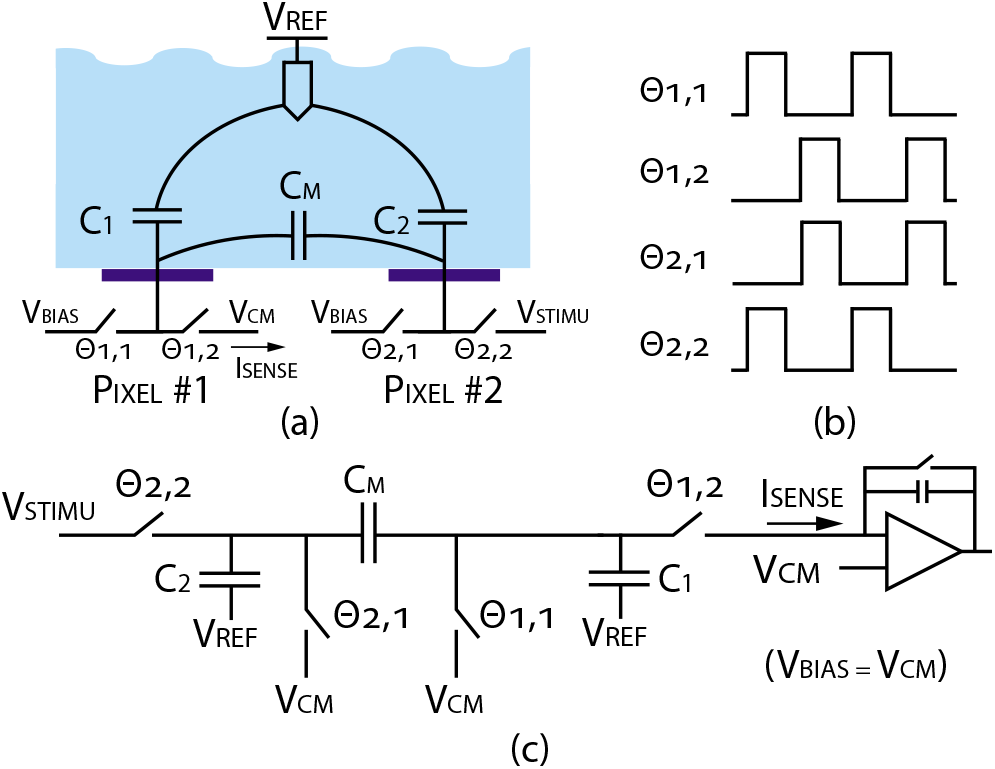
Illustration of the two-phase mutual capacitance measurement. (a) Two adjacent electrodes with a simplified capacitance model. (b) The pixel switches are driven by two sets of non-overlapping clocks. (c) If *V_BIAS_* = *V_CM_* and the clocks have 180° phase offset, the circuit becomes equivalent to a parasitic-insensitive switched capacitor integrator, which measures *C_M_*.

The electrodes are contained within pixels which can switch their bias voltages between multiple sources. The signal current *I_SENSE_* from electrode #1 is routed to a column amplifier where it is integrated and measured. *V_BIAS_* and *V_STIMU_* are provided from external voltage references, and *V_CM_* is the virtual ground potential of the current integrator. The timing diagram of the two sets of non-overlapping clocks is shown in Fig. 1(b), where Θ_1_ and Θ_2_ are 180° out of phase.

Interestingly, if we set *V_BIAS_* = *V_CM_*, this circuit can be equivalent to a classical non-inverting switched capacitor integrator as shown in Fig. 1(c). When both Θ_1,2_ and Θ_2,1_ are high, the voltage across *C_M_* is *V_STIMU_* − *V_CM_*. When both Θ_1,1_ and Θ_2,2_ are high, *C_M_* is discharged to 0V. The average integrated current can be expressed as *I_SENSE_* = *C_M_* (*V_STIMU_* − *V_CM_*) *f_clk_*. Since the voltage across *C*_1_ does not change, *I_SENSE_* is only a function of *C_M_*.

Fig. 2(a) illustrates the process of scanning the sensor array to form one impedance image, and then varying the scanned pattern to create multiple different impedance perspectives. In this simplified example, we use a 3 × 3 kernel, where the grid indices of Pixel #1 and Pixel #2 from Fig. 1(a) are related by (*i*_2_, *j*_2_)=(*i*_1_ + *δ_i_*, *j*_1_ + *δ_j_*), with *δ_i_* = *δ_j_* = 1. This kernel is scanned over the entire array to generate an image. The acquisition is repeated for different offset vectors (*δ_i_, δ_j_*), producing a collection of images with slightly different dependence on the sample’s spatially varying impedance. To acquire all pairwise N × N kernels requires measuring N^2^-1 images.

**Fig. 2:**
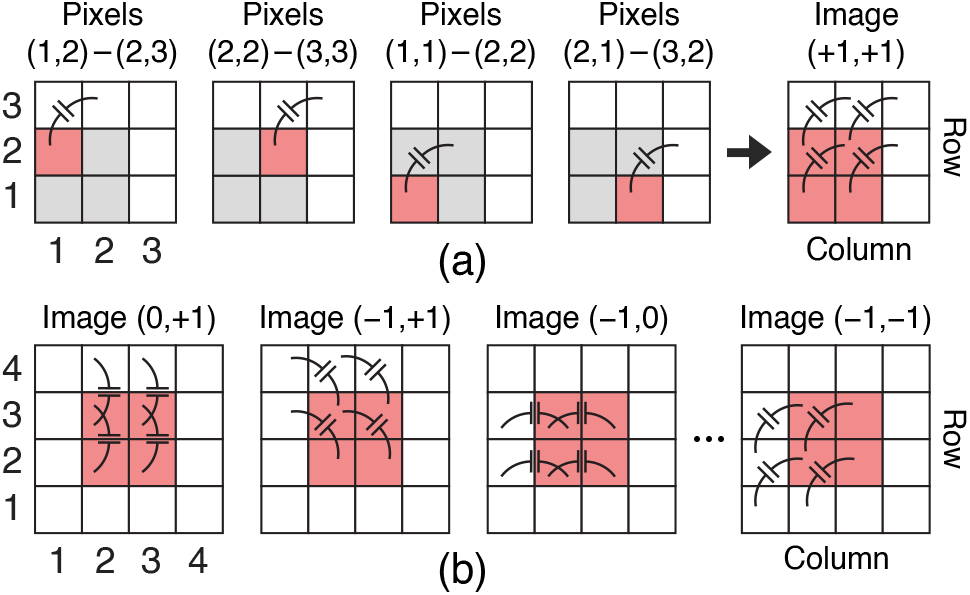
Illustration of mutual capacitance imaging. (a) Constructing an image from pairwise capacitance measurements, for a kernel offset of (*δ_i_, δ_j_*)=(+1,+1). (b) With a 3 × 3 offset kernel, 8 different kernel images can be captured, each giving a different perspective of the permittivity distribution above the array. Each image is labeled with its offset vector.

## III. CMOS Sensor Array Design

### A. Pixel and Array

Simplified schematics are shown in Fig. 3(a). The active sensing area has 10,000 pixels arranged in a 100 × 100 array. Each pixel can be driven by one of two pairs of non-overlapping input clocks (Θ_1,1/2_ and Θ_2,1/2_), whose relative phases are synthesized by an external delay lock loop. A set of control signals drive each row (R_*A/B*_) and column (C_*A/B*_), to determine the clock selection in each pixel. All switches are implemented as single NMOS transistors. After each pixel measurement, the NMOS switch gates are fully discharged to prevent stored charge from interfering with the next pixel scan. The output current of each pixel can either be routed to the readout circuit or to the *V_STIMU_* voltage reference as shown in Fig. 1.

**Fig. 3:**
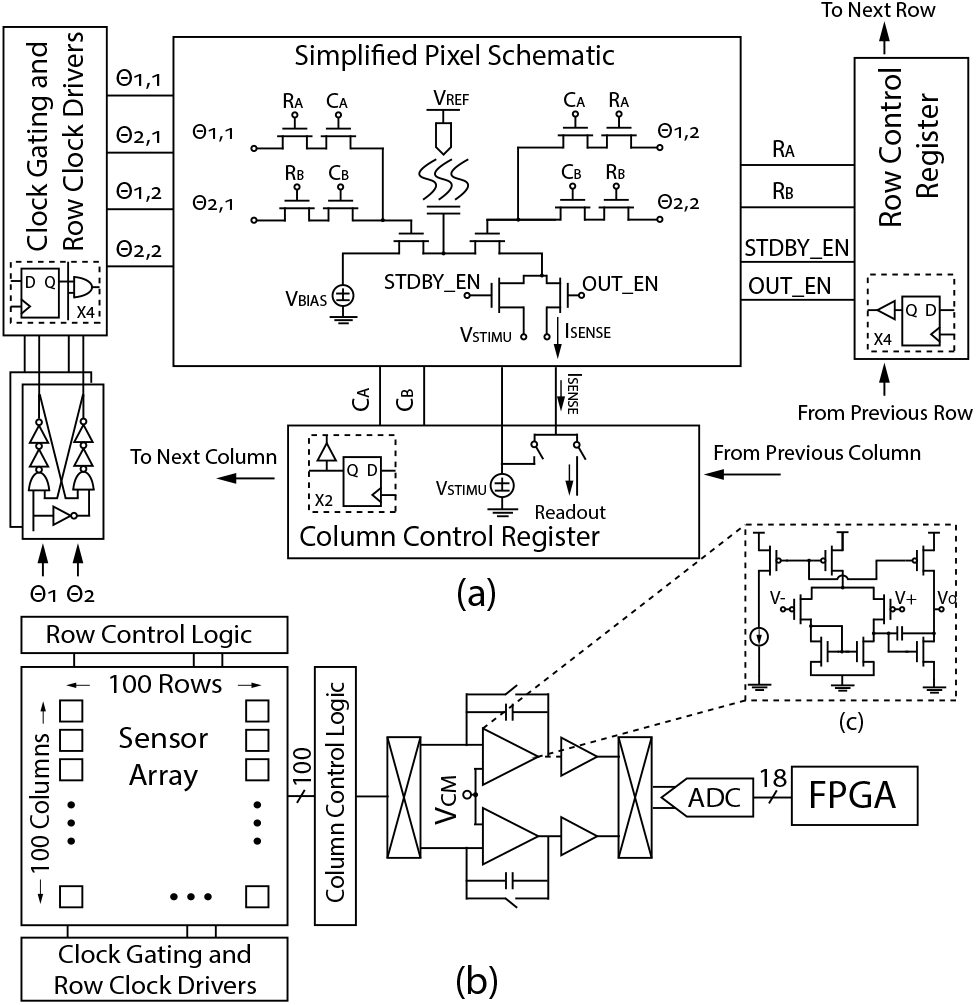
(a) Simplified pixel and array schematic. (b) Architecture of the 100 × 100 pixel sensor array, including the column readout signal path.

### B. Readout Circuit

The readout circuit shown in Fig. 3(b) includes chopping switches and a pair of integrators for signal amplification, followed by buffers that drive an external 500 kS/s 18-bit ADC. Differential chopping and correlated double-sampling are used to suppress the 1/f noise and offsets.

## IV. Experimental Results

### A. Layout and Packaging

The circuit is implemented in a 180 nm 1P6M CMOS process, occupying 2.24 mm^2^ (Fig. 4(a)). After encapsulating the wirebonds in epoxy, a simple open-top fluidic cell is assembled around the sensor (Fig. 4(b)) and the aluminum top metal is chemically removed to expose a titanium nitride electrode surface as previously described [9], [10].

**Fig. 4:**
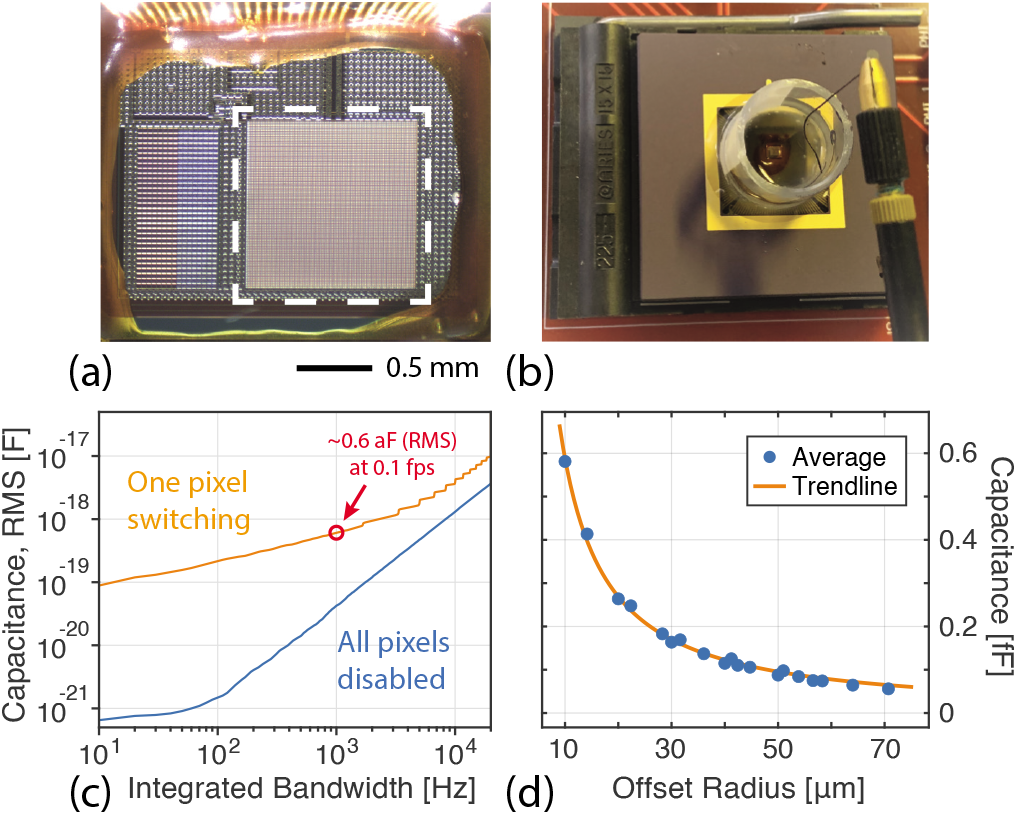
(a) Micrograph of the 100 × 100 sensor array. The active area is outlined in white. (b) Photograph of the packaged chip with a fluid chamber and silver/silver-chloride reference electrode. (c) Experimental RMS capacitance measurement noise for a switching frequency of 100 MHz, as a function of the integrator signal bandwidth. The integration bandwidth dictates the maximum image acquisition frame rate. (d) The mutual capacitance decreases with distance. In one dataset, the average measured mutual capacitance across the whole array e irically fit a power law, *C_M_* ≈ *ar^b^* ≈ 7.92*r*^−1.13^ fF, where 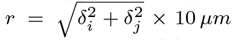. This average capacitance describes the bulk dielectric response of the water, but we would expect a similar trend in the Δ*C* signal that identifies algae cells.

### B. Sensor Array Characterization

During the measurement, the row and column addresses increment every 100 *μs*, and the chopping period and integrator reset interval are both 50 *μs*. The integration capacitance is 5 pF, and two output values are averaged for each pixel. At *f_clk_* = 100 MHz and *V_CM_* − *V_BIAS_* = 50 mV, the effective input-referred capacitance noise floor is shown in Fig. 4(c) as a function of the measurement bandwidth. The primary noise contribution is the kTC noise of the switched pixel capacitance, and at 1 kHz bandwidth (1 ms/address), we measure a noise floor of 0.6 attofarads (rms). We commonly allocate 0.1 – 1 ms/address. The total power consumption is 24.5 mW.

### C. Imaging Unicellular Algae

With their varied shapes and sizes, algae cells can be useful test cases for impedance imaging [11]. Here we selected *Cosmarium turpinii* for its intermediate cell size (approximately 50 *μm* diameter), as well as its notable bilobal shape (Fig. 5(d)), which is a useful starting point for resolving sub-cellular structure. In previous work [11], an earlier impedance sensor was able to image single cosmarium cells, but it could not resolve sub-cellular features.

**Fig. 5:**
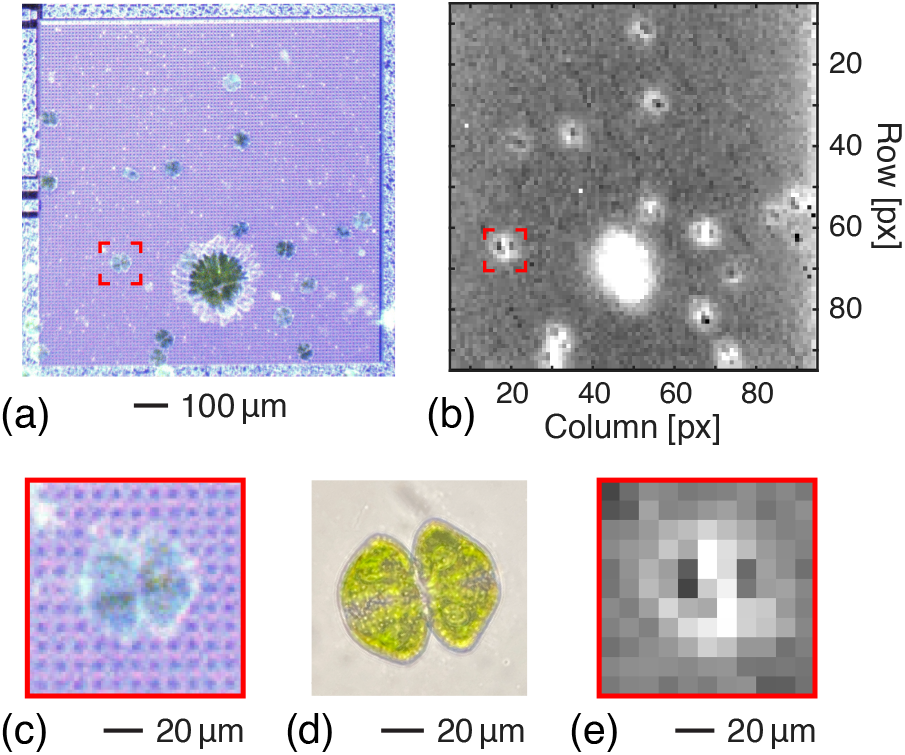
(a) An overhead optical inspection microscope image of the sensor, submerged in a solution containing green algae cells. The many smaller cells are *Cosmarium* and the larger one is *Micrasterias*. (b) A single impedance image for the kernel offset (+1,+2), with a switching frequency of 12.5 MHz. (c) A magnified region of the optical image showing a single algae cell on the CMOS electrode array. (d) A transmitted light bright-field microscope image of the same algae species (*Cosmarium turpinii*). (e) An 11 × 11 pixel area of the impedance image from panel (b) centered on the same algae cell whose optical image is shown in panel (c).

Fig. 5(a) presents an optical image of single algae cells on the surface of the sensor, and Fig. 5(b) shows an impedance image of the same sample. In a single image, we can sometimes coarsely detect the two hemispherical lobes of the algae cells. Fig. 5(c) and Fig. 5(e) zoom in to one cell.

In Fig. 6, we focus on one *Cosmarium* cell across 120 different mutual capacitance kernels, with the relative offset of Pixel #2 swept through an 11 × 11 pixel block. Each offset image provides slightly different amounts of contrast and detail about the cell, and has a distinct shift and stretch factor.

**Fig. 6:**
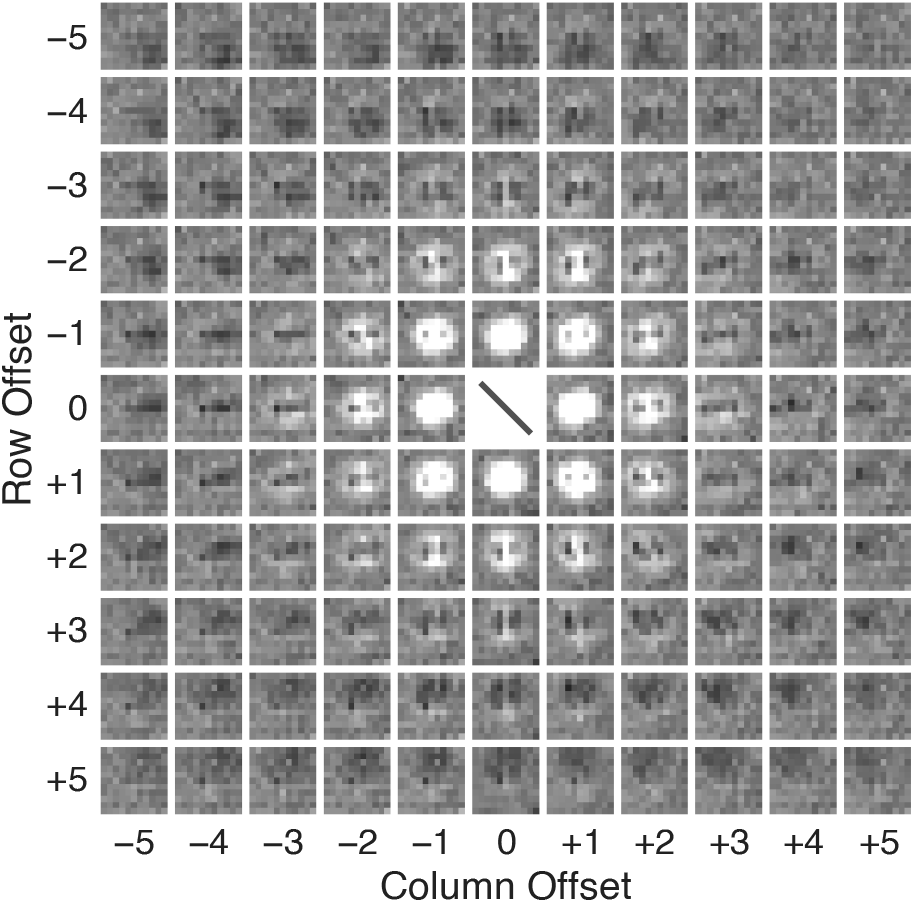
A montage of 120 different impedance images, using kernels with pixel offsets *δ_i/j_* ∈ [−5, 5]. The images are cropped around an 11 × 11 pixel region containing the same algae cell from Fig. 5(c). The color scale of each image has been adjusted independently (Tukey fences with a sensitivity of 3) to improve feature contrast for visual comparison.

### D. Super-resolution Impedance Imaging

Many techniques have been developed for assembling a high-resolution optical image from multiple lower-resolution images. For example, multiple video frames can be aligned and computationally merged, taking advantage of the fact that camera motion produces subtle spatial shifts in the scene relative to the image sensor. Thanks to sub-pixel shifts in the scene, combining multiple re-aligned video frames can yield a composite image with higher resolution than the individual frames [8].

Similarly, impedance images recorded using different kernels have slightly different effective spatial sampling, as we saw in Fig. 6. The mutual capacitance between two pixels is a complex function of the 3-D electric field distribution within the sample, but as a starting point, we can consider each image to represent the average permittivity of the sample in the space between Pixel #1 and Pixel #2. While this necessarily implies some loss of resolution to spatial averaging, it also adds signal diversity between frames which can enhance digital reconstruction.

To produce a composite super-resolution impedance image, we upsampled the original images by 16× using a Lanczos-3 kernel, shifted them by (−*δ_i_*, −*δ_j_*), and performed a shear mapping with 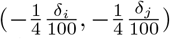 to partially compensate for the offset-dependent image skew. The stack of re-aligned images was then summed together. As one might anticipate, averaging multiple measurements reduces uncorrelated background noise, and the fractional differences in image grid registration allows for finer foreground features to be enhanced. Fig. 7 shows a composite image constructed from the 120 kernel offsets shown in Fig. 6. Since the mutual capacitance reduces super-linearly with distance (Fig. 4(d)), there can be diminishing returns from including images with far-spaced electrode offsets. This super-resolution image is a first proof of concept, and there are many opportunities for improved algorithms to align and merge multiple impedance images. Different kernel offsets emphasize different features within the algae cells (Fig. 8), and it would be natural to imagine combining these types of measurements with algorithms from electrical impedance tomography [12] to generate a 3-D reconstruction of the sample permittivity.

**Fig. 7:**
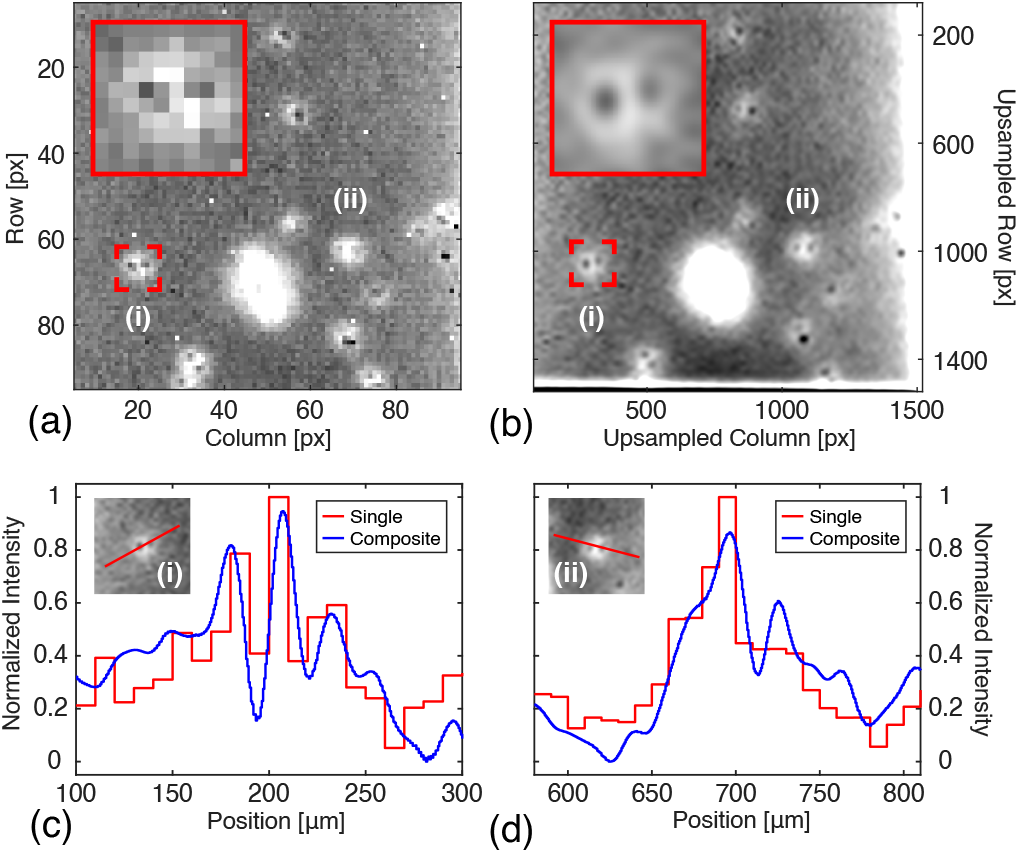
Super-resolution impedance imaging of algae cells. (a) A single impedance image acquired with (*δ_i_, δ_j_*)=(−2,−1). The red box highlights the position of the magnified inset. (b) A composite impedance image made by upsampling (16×), shifting (by the kernel offset vector), shearing (by half the offset over twice the array size), and finally summing the impedance images from the 120 kernel offsets shown in Fig. 6. (c,d) Slices of the impedance image taken across the isthmus of two cosmarium cells to highlight their bi-lobal structure. The sub-cellular features are sometimes visible in single impedance images and are enhanced in the composite data.

**Fig. 8:**
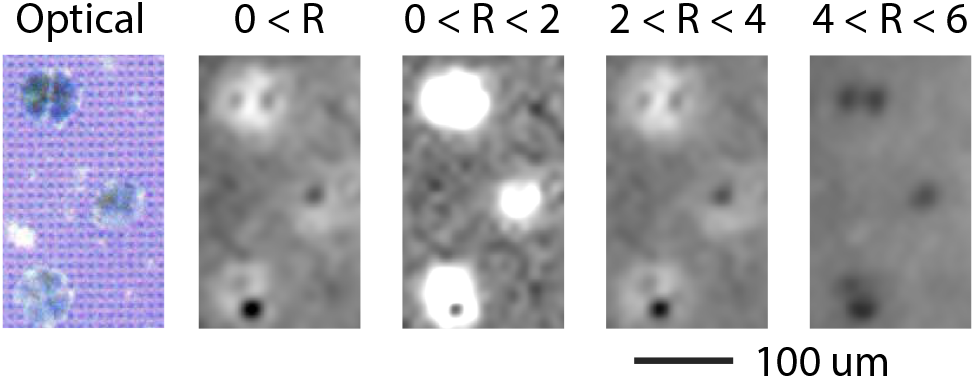
Kernels with longer offset vectors highlight different features in the scene. By constraining the composite image reconstruction to different offset vector magnitudes 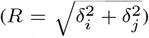, we can see that shorter pixel separations detect the whole *Cosmarium* cell, while larger pixel separations have better contrast for the pyrenoid in each semi-cell.

## V. Conclusion

We have presented a new approach for spatially-resolved electrochemical impedance imaging, built around a CMOS sensor designed with a flexible and area-efficient strategy for pairwise capacitance measurements. In addition to improved sensitivity compared to previous platforms, the system presents a new path for impedance sensing to take advantage of computational strategies from optical imaging. As a first demonstration, we presented state-of-the-art non-optical measurements of unicellular green algae, producing super-resolution impedance images which resolve subcellular spatial structure in electrical permittivity. There are opportunities to improve on these results with more advanced algorithms, and a wide array of biomedical applications stand to benefit from spatially resolved electrochemical imaging.

## Supplementary Methods

### 1. Instrumentation

Electrochemical impedance images were acquired using the custom CMOS sensor presented in the main text. The resulting images were processed and analyzed with custom MATLAB scripts. Images of the sensor (Fig. 4) were taken with a stereo microscope (Olympus SZ61TR, 3× magnification). A reference optical image of an algal cell (Fig. 5d) was taken using a Nikon TI-U inverted microscope (bright-field, 20× magnification). Optical images of the cells on the CMOS sensor (Fig. 5a & c, Fig. 8 left panel) were taken using an inspection microscope (Edmund Optics #55-150 dual tube body, 0.75 – 3× magnification, FLIR BFS-U3-51S5C-C camera).

### 2. Living Specimen

Live algae were purchased from Carolina Biological Supply Company (Burlington, NC, USA). The *Cosmarium* (Item #152140) and *Micrasterias* (Item #152345) cells were obtained as unialgal cultures.

### 3. Experimental Setup

Before each experiment, a small fluid chamber (Fig. 2b) was fashioned out of a section from a 10 mL centrifuge tube and bonded to the surface of the sensor using silicone elastomer. The chamber is then filled with aluminum etchant (Type A, Transene Company, Danvers, MA, USA), to remove the aluminum top metal of the sensor chip [1]–[3]. After etching the electrodes, chamber was thoroughly cleaned with DI water, and filled with phosphate-buffered saline (0.1 M potassium phosphate, 1 M potassium chloride). An Ag/AgCl wire was placed into the solution to serve as a reference electrode. The entire setup was positioned under an inspection microscope and algae samples are pipetted into the chamber. After the cells settle onto the sensor surface, optical images are taken for reference and electrochemical imaging is performed. Each 100 × 100 impedance image takes approximately one second to acquire. Data was acquired through an FPGA module (XEM6310-LX45, Opal Kelly) over a USB 3.0 interface.

